# Unmasking complex kinetics in viral entry by inferring hypoexponential models

**DOI:** 10.1101/2025.05.18.654751

**Authors:** Oyinkansola Adenekan, Peter M. Kasson

## Abstract

Single-event completion times, such as are estimated in viral entry, offer both promise and challenge to kinetic interpretation. The promise is that they are able to constrain underlying kinetic models much more efficiently than bulk kinetics, but the challenge is that completion times alone can incompletely determine complex reaction topologies. Gamma distributions or mechanistic models have often been used to estimate kinetic parameters for such data, but the gamma distribution relies on homogenous processes contributing to the rate-limiting behavior of the system. Here, we introduce hypoexponential analysis to estimate heterogeneous kinetic processes. We demonstrate that hypoexponential fitting can indeed estimate rate constants separated by 2-3 orders of magnitude. We then apply this approach to measurements of SARS-CoV-2 entry, showing that ACE2 reduces the number of rate-limiting steps but does not change the rates of these kinetic processes. We propose a kinetic model whereby SARS-CoV-2 entry is driven by a mixture of ACE2-accelerated and ACE2-independent spike protein activation events. Inferring such models requires the capability to detect heterogeneous kinetic processes, provided by robust estimation of hypoexponential distributions.

## Introduction

Biophysical processes pass through one or more transitions or energetic barriers. Single molecule experiments measure fluctuations in these biophysical processes, giving potential to understand detailed mechanistic information in a way that traditional experimental techniques that average over molecules obscures. One single molecule experimental output is several measurements of time to reaction completion or dwell times (1). From this distribution of dwell times, we can better understand the molecules’ behavior by identifying a sequence of conformationally stable states and quantifying transitions between these states(2). But different underlying states can have the same experimental signal/output. Therefore, recovering the correct number of underlying states and the transitions between them remains an analysis challenge. In particular, we wish to use reaction kinetics to better understand how SARS-CoV-2 and influenza enters cells. We conducted single-molecule fluorescence experiments to measure how long it takes for these virus membranes to fuse with model host membranes. Extracting kinetic parameters from these dwell times can facilitate development of mechanistic models of viral entry.

There exist several methods for quantifying kinetic states from experimental time domain data. Hidden Markov models (HMMs) (3-7) have long been applied for this purpose, and both HMMs and continuous time Markov models have been used to identify transitions between states from ion channel patch clamp experiments (8). Similarly, different HMM have been used to extract kinetic information from smFRET experiments (2,9-11). HMMs are a useful tool for analyzing experimental data where, in general, kinetic states have distinct experimental emissions. Here, we address a different problem: single-event kinetics where the readout is effectively a step function: θ(s-s_N_) such that the start state s_0_ and all intermediate states s_N.._s_N-1_ have emission 0 and the product s_N_ has emission 1. HMMs tend to be highly degenerate for this class of problems, driving the development of other kinetic fitting approaches.

Previous studies fit dwell times from single-molecule fluorescence experimental data to a gamma function to extract reaction kinetic parameters (12,13). A gamma function describes a multi-step, linear process. Researchers have used gamma analysis to develop a model of influenza entry to host cells where each kinetic step is an independent binding event between the virus’s spike protein and the protein HA (12). However, since a dwell time distribution’s shape is mostly determined by the reaction’s slowest rate, the gamma distribution assumes that each transition’s rate constant is equal to the slowest rate. By assuming that each step in a reaction is equal to the slowest step, the gamma distribution potentially obscures the kinetics of a reaction, making it a limited tool for analyzing dwell time distributions. Cellular automaton models are another method for extracting kinetic information from dwell time distributions and have also been used for analysis of viral entry data from single-event experiments (14-17). These models require more detailed mechanistic knowledge of a reaction beyond the dwell time distribution and are thus not considered here.

In this paper, we propose recovering reaction kinetics by fitting dwell times to the hypoexponential distribution. The hypoexponential distribution is in the same family of distributions as the gamma, but it estimates a different rate for each step in a sequential process. This distribution may be well suited for estimating both number of transitions and unique rates of transition for a multi-step reaction process. However, the hypoexponential distribution may have identifiability issues: as the number of states increases, the harder it becomes to estimate this parameter accurately (18,19). Despite this potential issue, the hypoexponential distribution may be a better analysis tool than the gamma for recovering kinetic information from dwell time distributions.

In this article, we will do the following 3 things: 1) quantify the difference in performance between gamma and hypoexponential for estimating reaction kinetics, 2) explore the quantitative limits of the hypoexponential distribution, 3) apply hypoexponential analysis to single molecule fluorescence data on influenza and SARS-CoV-2 entry. We will show that for 2 and 3 step processes, a hypoexponential distribution more accurately recovers reaction kinetics in synthetic experiments. Additionally, we will explore the difficulty with estimating kinetics as the number of steps increases. Finally, we will apply hypoexponential analysis to dwell times collected via single-molecule fluorescence experiments for both SARS-CoV-2 entry and influenza entry to host cells.

We will demonstrate that while the hypoexponential distribution has limitations for reactions with higher numbers of steps, it does provide significant benefits over the gamma distribution in terms of information gained.

### Theory

We extract kinetics of biophysical reactions by fitting chemical reaction models to observed single-event reaction dwell times. One way to generate such reaction models is to decompose a complex reaction into a network where each edge represents a quasi-first-order reaction. The cumulative distribution function for each edge is thus an exponential function parameterized by a rate *k*. Here, we consider linear processes, as shown in Equation 1.

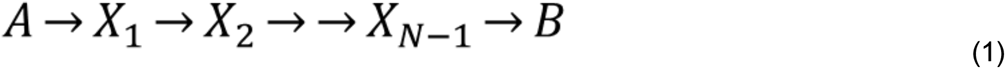

The biophysical rationale for this approach is that a single-barrier chemical process will display exponential kinetics of this form according to Eyring’s law. A sequential process, which models a multi-step biophysical reaction, thus becomes convolution of *N* exponential random variable. A handful of distributions arise from convolution and other aggregations of 2 exponential random variables. Here, we focus on the gamma and hypoexponential distributions.

Previous studies fit reaction dwell times to the gamma distribution to extract kinetics. The gamma distribution is the convolution of *N* exponential random variables, each with identical rate parameter *k* (Equation 3). Thus, the gamma is parametrized by *N*, the number of transitions in the process, and *k* the rate of completion for any single step. In a biophysical reaction, *N* corresponds to the number of free-energy barriers that contribute rate-limiting steps in the reaction. Notably, the gamma distribution assumes that *k* is the same for each step. Because the slowest step contributes most to the shape of the dwell time distribution, *k* typically reflects the slowest step in a reaction <ref>.

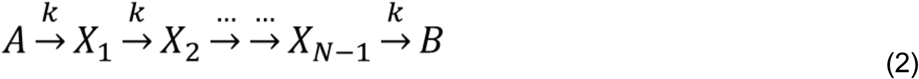

In this paper, we fit hypoexponential functions to observed reaction dwell times. Like the gamma function, a hypoexponential is the convolution of several exponential random variables. But in a hypoexponential, the number of exponential random variables convolved is specified rather than inferred. Equation 3 is a 2-parameter hypoexponential parameterized by *k*_*1*_, the rate of the first transition, and *k*_*2*_, the rate of the second transition. Unlike the gamma function, a hypoexponential function has an independent rate for each transition in the modeled process. But because the hypoexponential specifies the number of transitions, we cannot directly estimate the most-likely number of transitions in a reaction by fitting a single distribution to the data. To address this, we estimate hypoexponential distributions with increasing numbers of transitions, *N*=1,2,…. In this case, estimation is performed using Metropolis-Hastings Monte Carlo as described below. Then we select N by identifying the model with the lowest corrected Akaike Information Criterion (AICc) (20,21).

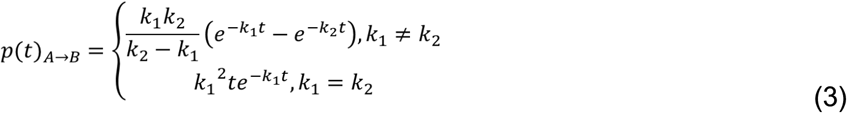

## Methods

### Single virus fluorescence experiments

Single-virus kinetics of influenza membrane fusion were performed according to previously published protocols (22) with the exception that the microfluidic flow cell was maintained at 32 ºC. X-31 (A/Aichi/1968 H3N2) influenza virus was bound to GD1a glycosphingolipid receptors displayed on liposomes immobilized in a microfluidic flow cell, and fusion was triggered by buffer exchange to pH 5.0. Fusion is assessed by dequenching of Texas Red-DHPC dye loaded into the viral envelope at a quenching concentration. Single-event dwell times were provided by Robert Rawle.

Single-virus kinetics of SARS-CoV-2 virus-like-particle fusion were previously reported (23), and the data are re-analyzed here. Kinetics were measured using fluorescently labeled pseudovirus fusing to liposomes, with attachment mediated by synthetic DNA-lipid attachment factors and fusion triggered using soluble protease. Experiments were performed with and without soluble ACE2 receptor. These data take the form of single-virus dwell times: the interval between fusion triggering of bound virus using a soluble protease and the fluorescence dequenching that serves as a real-time fusion signal.

### Data simulation and modeling

For initial validation, we generated synthetic data corresponding to known ground-truth values of *N* and *k*_*i*_ using the Gillespie algorithm(24,25) implemented in the *pysb* and *bionetgen* Python packages and APIs. We simulated 300 dwell times for each set of ground-truth values. Plotted data were generated using a 1 s granularity, matching SARS-CoV-2 experimental parameters. We also performed sensitivity analyses with 1 ms granularity.

Fitting of hypoexeponential distributions to both synthetic and experimental data was performed using a simple Metropolis-Hastings algorithm written in Python. The algorithm proposes a set of values for the rate parameters and then accepts or rejects the proposal based on its likelihood given the observed data. Code can be found on Github: https://github.com/kassonlab/hypoexponential-analysis. Gamma distributions were fit to the data using previously published code (26).

### Evaluating hypoexponential and gamma fits

We evaluated the performance of both hypoexponential and gamma fits (k and N) using percent error: error_k_ = abs(k_ground truth_-k_estimated_)/k_ground truth_*100 and error_N_ = abs(N_ground truth_-N_estimated_)/N_ground truth_*100.

## Results

### Extracting kinetic parameters on viral entry process by fitting data from single virus fluorescence experiments to reaction models

We briefly describe the experimental design and analytical goals, followed by validation on synthetic data and application to problems in viral entry. We use single-virus optical microscopy where fusion kinetics are measured using one of two readouts: either a fluorescent reporter of lipid mixing or a fluorescent reporter of content mixing. In both cases, the unfused virus and all intermediates until the process of interest occurs have baseline fluorescence, and the fused conjugate has elevated fluorescence. The approach is schematized in Figure 1, and details on the experimental protocol have been given in a recent methods paper (27). Briefly, target liposomes are immobilized in a flow cell, and viral particles are added and bound to them. Binding occurs either via endogenous receptors or synthetic DNA tethers (26). Each viral particle that fuses generates an abrupt increase in fluorescence, which can be extracted as the waiting time to fusion. Collating these waiting times from many individual particles yield a distribution of dwell times that can be represented using an empirical cumulative density function (eCDF). We model this curve as a biophysical reaction that starts in an unfused state and traverses through a number of intermediate states N before reaching the final fused state. We fit dwell times from single virus fluorescence experiments to the aforementioned reaction model to get estimates of N and rate constants *k*_i_ for transitions between states.

**Figure 1.**
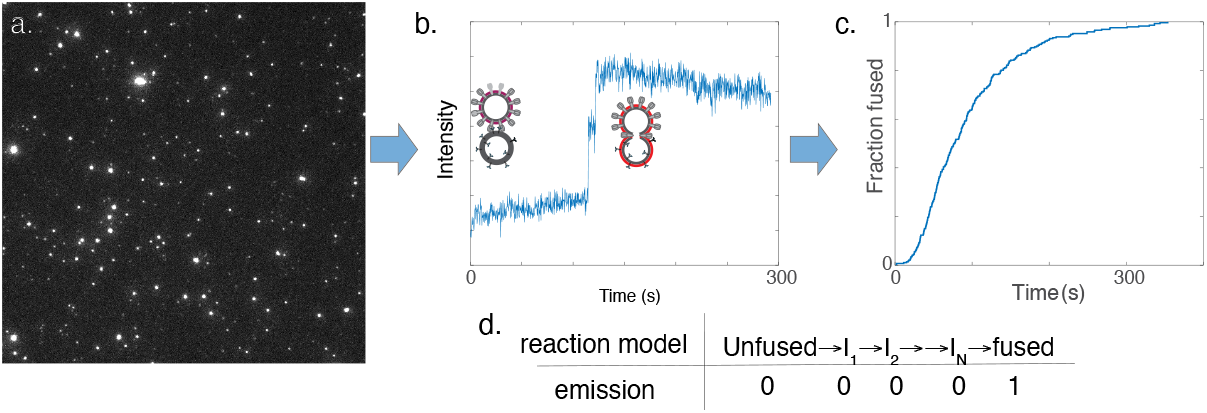
Representative data showing a still from microscopy of influenza virus undergoing fusion (a), a single-virus intensity trace with schematized unfused and fused membranes before and after the intensity jump (b) , the CDF compiled from these traces (c), and a linear reaction model with corresponding 0 and 1 emissions denoting pre- and post-jump intensity traces (d).

### Hypoexponential fitting outperforms gamma analysis in reconstructing generative reaction models for synthetic data

In evaluating the utility of hypoexponential fitting, we first asked whether it could outperform gamma analysis when the ground-truth reaction model was known. To that end, we created synthetic data using reaction models consisting of 2 or 3 kinetic steps and where the k_i_ values varied from identical to 100,000-fold different. Rate values were chosen to cover the ranges anticipated for influenza and SARS-CoV-2 single-virus fusion kinetics. Synthetic data was created by sampling dwell times using the Gillespie algorithm(24). We then compared the results of hypoexponential fitting and gamma analysis on these synthetic data. Figures 2 shows eCDFs for three synthetic data examples and the corresponding gamma and hypoexponential fits. In general, both gamma and hypoexponential distributions produced near-equivalent fits when the ground-truth rate constants were identical or when they were so different that the fast step is no longer rate-limiting. Both overall fit the CDFs through the entire range, but subtle differences were appreciable when the fast and slow rates differed by a moderate amount. In subsequent figures, we compare gamma and hypoexponential performance on individual rate estimates.

**Figure 2.**
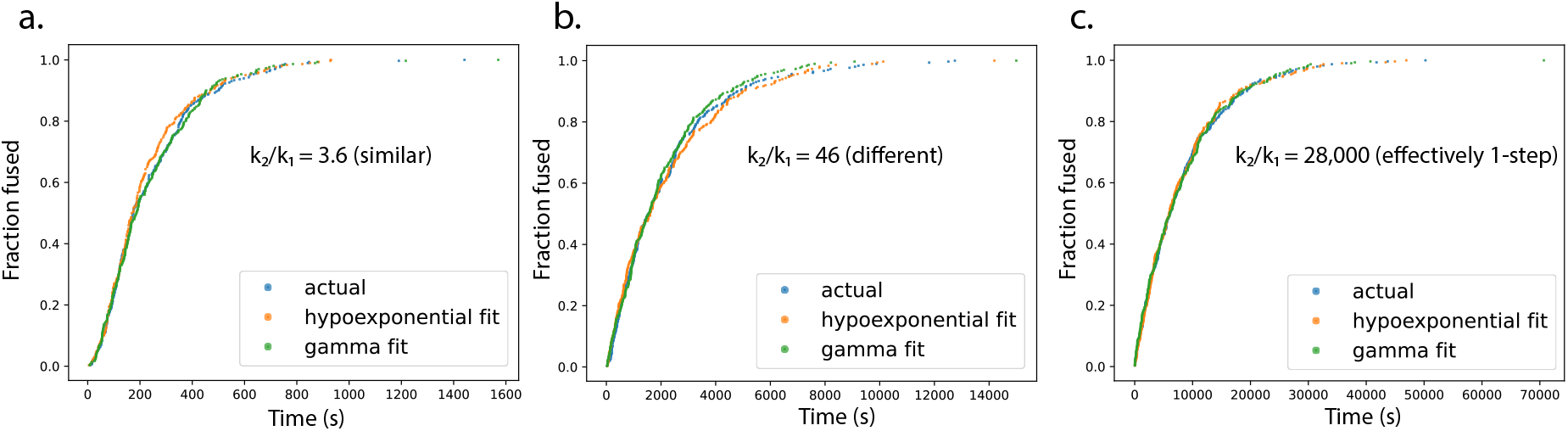
Cumulative distribution functions and fits plotted for three scenarios, where k_1_ and k_2_ are similar (a), where they differ substantially (b), and where they differ sufficiently that k_2_ is no longer rate-limiting. Rate constants used were 5.92e-3 and 2.13e-2 s^-1^ (a), 4.59e-4 and 2.13e-2 s^-1^ (b), and 1.28e-4 and 3.55 s^-1^ (c).

**Figure 3.**
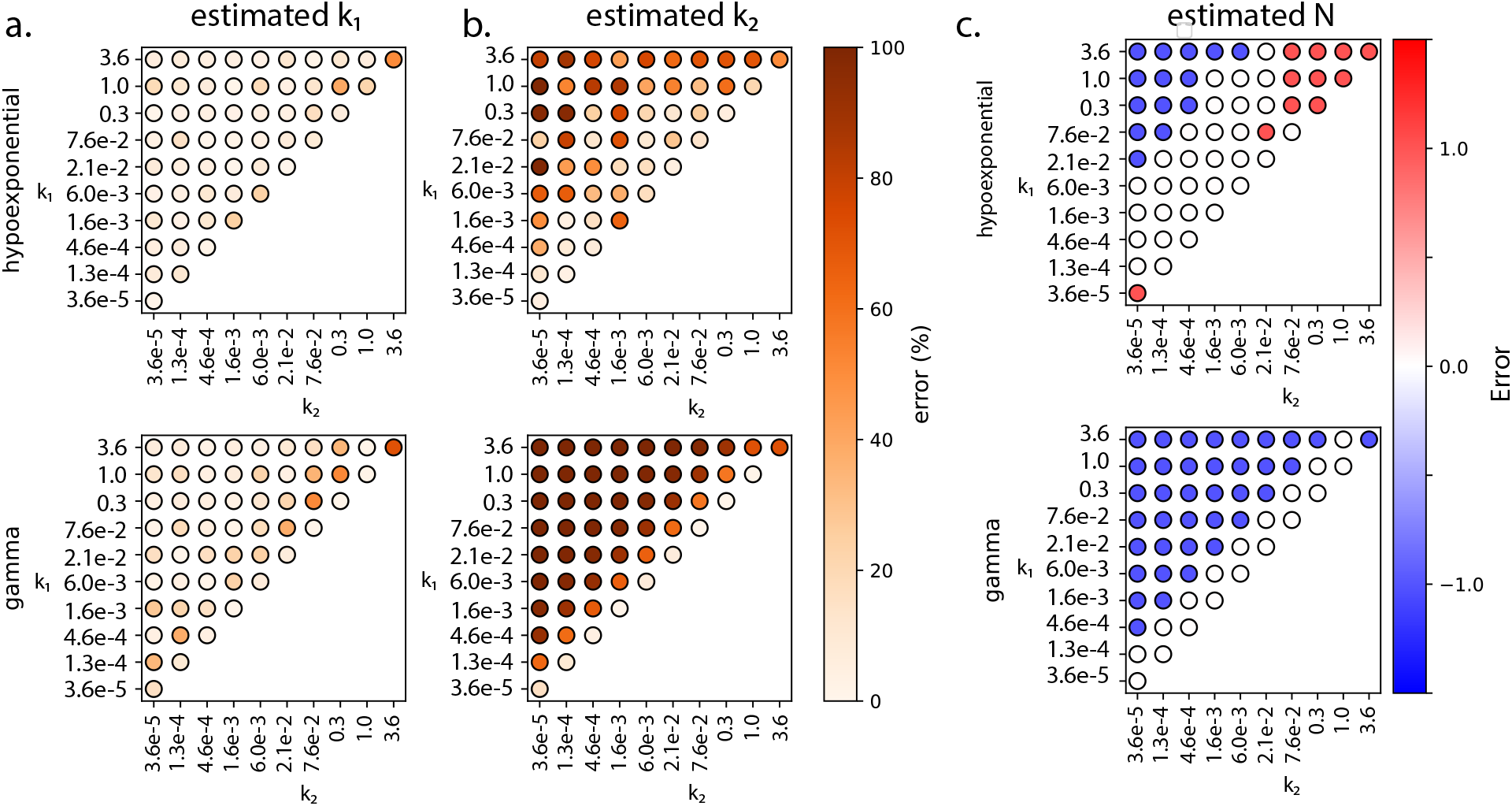
Performance comparison of gamma and hypoexponential fits for two-step reactions. Synthetic data from a range of k_1_ and k_2_ values were fit using either gamma or hypoexponential MCMC approaches. Percent error in estimated k_1_ values (a), k_2_ values (b), and absolute error in estimated number of steps N are plotted. Hypoexponential fits outperform gamma in all three recovery tasks except when the two rates are equal.

### Performance on two-step reactions

As shown in Figure 2, hypoexponential fitting better recovers both rates and number of steps in the ground-truth reaction model than does gamma analysis. Interestingly, for the slower step, hypoexponential and gamma fitting recover rates with similar accuracy. Hypoexponential fitting recovers about 90% of rates with less than 20% error, while gamma fitting recovers about 70% of the slower rates at the same error level. For the faster step, however, hypoexponential fitting substantially outperformed gamma analysis. While both methods display increased error as rates deviate from one another, hypoexponential fitting is less affected by this. For the hypoexponential fitting, percent error is consistently above 50% when rates are more than 100-fold apart. For gamma recovery of the slower step, percent error is always above 50% when rates are not identical. This highlights the weakness of gamma fitting: because its underlying model assumes identical rates, it is fundamentally unable to recover rate constants that differ substantially from each other.

Recovery of the number of steps (N) displays a similar trend: worse recovery as the rates deviate more but better performance with hypoexponential fitting than gamma. Hypoexponential fitting generally recovers N accurately for rates differing less than 1000-fold, while gamma fitting only recovers N when rates are within 100-fold. Interestingly, hypoexponential fitting seems to struggle to recover Ns for pairs with similar rates that are extremely fast or slow. Overall, hypoexponential fitting outperforms gamma fitting in recovering rates and number of kinetic steps (55% vs. 33%) in 2 step synthetic reaction schemes. These synthetic experiments establish hypoexponential fitting as a useful tool for extracting kinetics, particularly when rates differ within 100-to 1000-fold.

### Performance on three-step reactions

We also tested hypoexponential fitting versus gamma analysis on 3-step synthetic reaction schemes, and it again displayed better recovery of kinetic parameters. For the slowest step, hypoexponential fitting slightly outperforms gamma analysis: recall of 89% to ≤20% error versus 63% to ≤20% error. Similar to the 2-step analysis, as the rates deviate from one another, hypoexponential recovers the middle rate much better than gamma: 58% to within 50% error, versus 25% to within 50% error. For the third and fastest step, hypoexponential fitting recovers 30% to ≤50% error, while gamma fails on all cases when the rates are non-identical. These results accentuate the trends from the 2-step data: both approaches are challenged by multi-step reactions with highly differing rates, but hypoexponential fitting performs better in these scenarios. This is in line with the gamma function’s underlying model of identical rates: it consistently recalls the slowest step in a reaction but is not flexible enough to estimate differing rates. Finally, hypoexponential fitting generally recovers N accurately when rates are within 100-fold of one another, whereas gamma only recovers N accurately when rates are within 10-fold, showing the same trend as the 2-step reactions but increased stringency due to the added kinetic complexity. While hypoexponential fitting recovers kinetics more accurately than gamma for 3 step kinetic systems, the reliability in recovering the fastest rate is low. As discussed later, this may reflect a fundamental statistical limit in reconstructing diverse rates.

For 2-step kinetic systems, the highest accuracy in recovering number of steps is achieved using the randomness parameter. This measure is derived from the relative variance of single-event waiting times and reports on the minimum number of steps in a linear scheme required to generate the observed waiting times (1,28). Not surprisingly, ceil(N_min_), where N_min_ = 1/*r*, the randomness parameter, performed better than either AICc on the hypoexponential fit or the gamma N parameter on our synthetic data test set (Fig. 4). Interestingly, the failures for this estimator occurred on a number of equal or near-equal rates, suggesting that an optimal estimator might account for some stochastic noise and take the form ceil(N_min_ - ϵ) for some tolerance hyperparameter ϵ. The rationale for this formulation is that if stochastic error is excluded, a N_min_ of 2.2 would require at least 3 steps, so ceil(N_min_) encodes this dependence. The ϵ is introduced to account for stochastic error, in effect differentiating N_min_ of 2.0± ϵ from N_min_ of 2.2.

**Figure 4.**
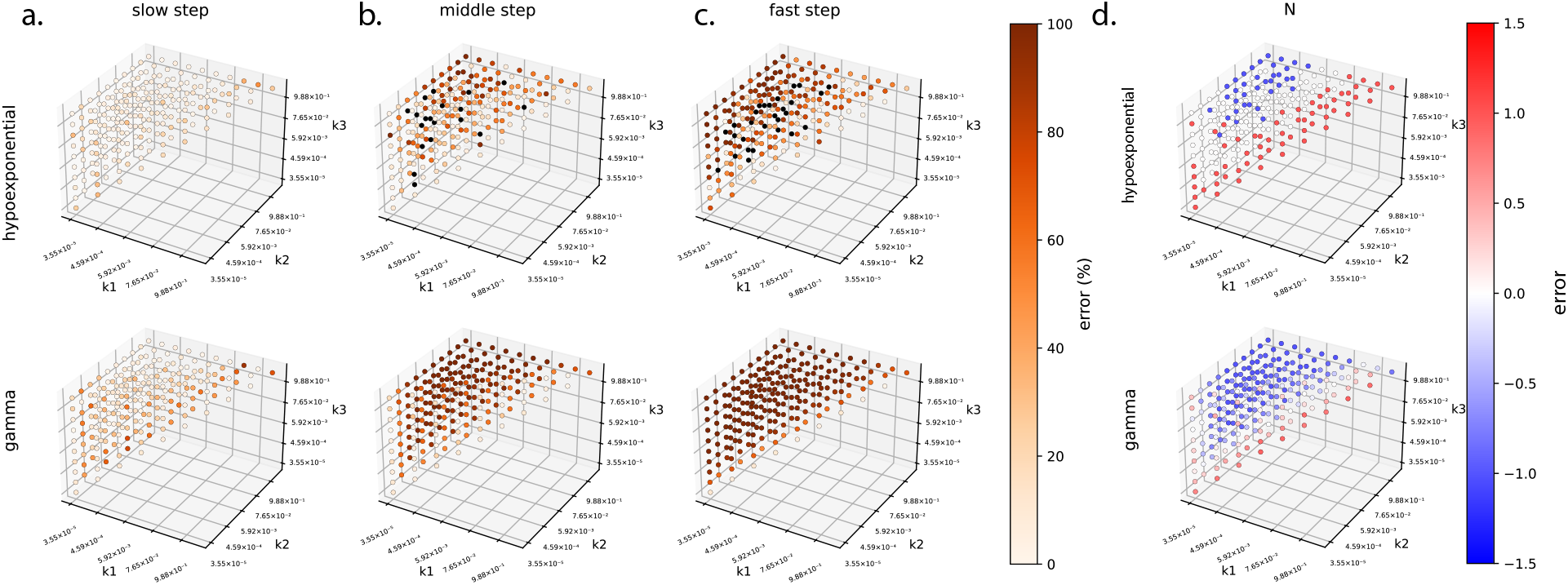
Performance of gamma and hypoexponential fits on synthetic data from three-step reactions. Percent error in recovery of the slowest, middle, and fastest rate constants is plotted in panels (a-c); absolute error in recovery of the number of steps is plotted in panel (d). Hypoexponential fits outperform gamma fits in cases where the rates are moderately different, substantially extending the range of accurate rate recovery.

**Figure 5.**
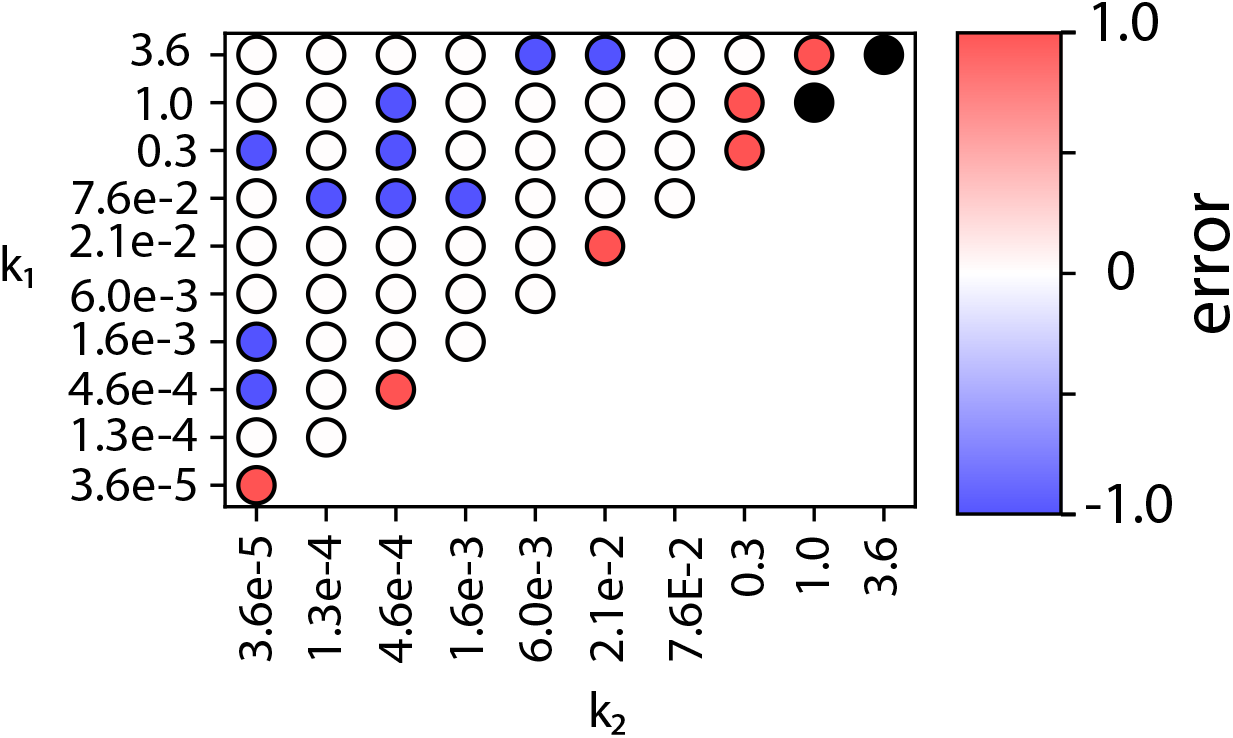
The randomness parameter provides a robust estimate of the number of steps in a heterogeneous kinetic process. Shown is the error in the number of steps using ceil(N_min_) for 2-step processes with varying rates. As can be seen, this estimator overall outperforms either AICc on hypoexponential fitting or gamma fitting but has some increased error along the “identical rate” diagonal. We therefore conclude that ceil(N_min_ - ϵ) for ϵ ∼ 0.1 is likely the optimal estimator for N_steps_, and this should be used to guide hyperparameter choice for hypoexponential fitting. Black points denote an error > 1.0, in this case produced by rates too fast for the 1s sampling interval.

### Estimating multi-step kinetics of influenza viral entry

Influenza virus undergoes a multi-step reaction to fuse its envelope with endosomal membranes in the cell and release the viral genome into the cytoplasm. This fusion reaction has been extensively studied using both bulk and single-virus kinetics experiments (12,14,26,29,30). Prior fitting of influenza fusion kinetics at the single-virus level has primarily utilized either gamma functions or more mechanistically detailed cellular automaton model (12,14,15,31); the randomness parameter has also been used as a means to determine the minimum number of sequential steps in the underlying kinetic scheme (32,33). Lipid mixing, or exchange of labeled lipids between the influenza virion and a target membrane, was modeled as 2-4 identical steps, with the greatest likelihood being 3 identical steps in most cases (12). Based on mutational experiments, the rate-limiting step was linked to pH-dependent release of the hemagglutinin fusion peptide (14).

We applied hypoexponential fitting to better understand the kinetic steps leading to influenza entry. The maximum-likelihood parameters for influenza lipid mixing using the data in Fig. 6 were two steps, which agrees well with gamma fits and randomness-parameter analysis on this dataset.

**Figure 6.**
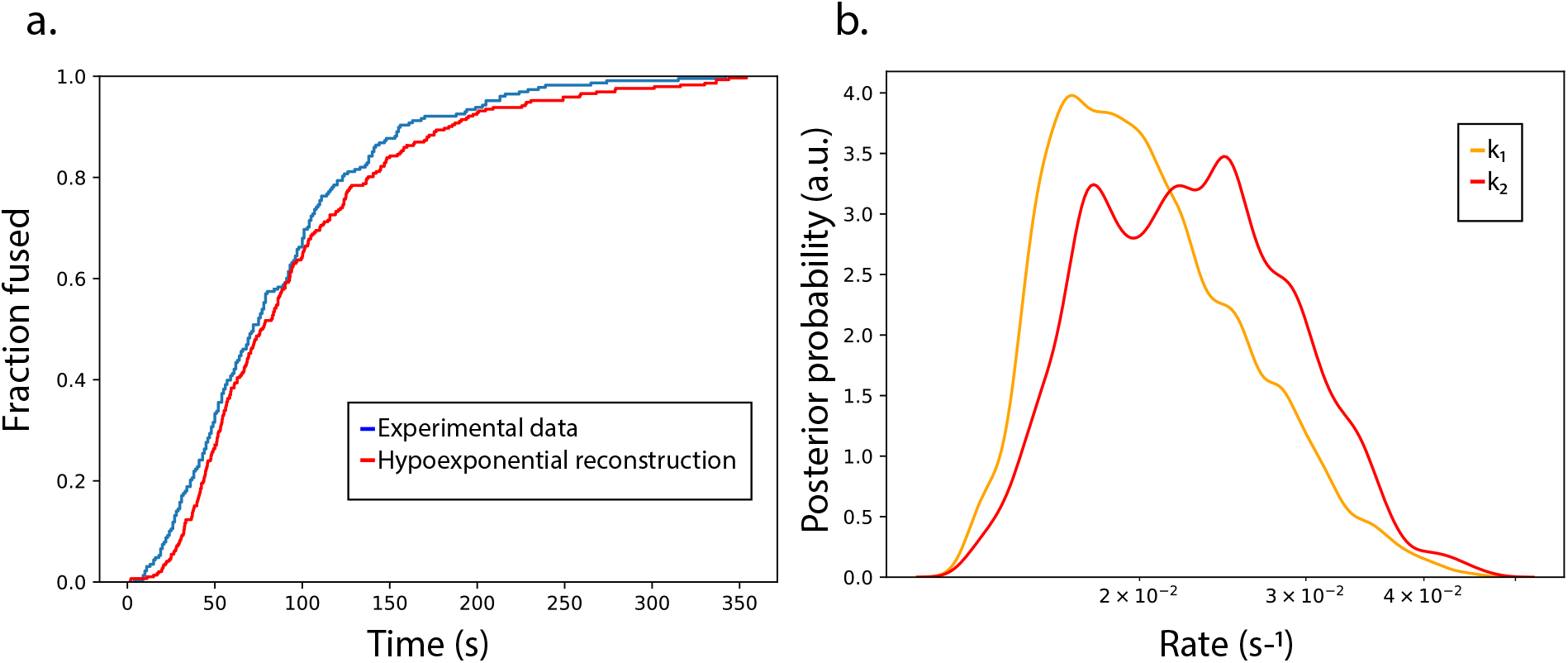
Hypoexponential fit to influenza lipid mixing data. Plotted in (a) are empirical CDFs for the observed lipid-mixing times (blue) and the hypoexponential fit (red). Plotted in (b) are posterior probability density estimates for k1 and k2 in the hypoexponential fit. Maximum-likelihood rate estimates are 0.022 s^-1^ and 0.020 s^-1^.

For a more detailed comparison, we leveraged the MCMC fitting process to examine posterior probability distributions and confidence intervals for rates rather than simple maximum likelihood estimates. Those are plotted in Fig. <N> to correspond to the maximum-likelihood estimates in Fig. <M>. For lipid-mixing data, posterior distributions corresponding to the number of kinetic steps predicted by the randomness parameter show N indistinguishable rate constants k_N_. Models corresponding to N+1 kinetic steps show N rates slightly slower than k_N_ and one rate much faster than k_N_. These results, as well as the synthetic data, demonstrate that hypoexponential fitting well captures two-step and typically three-step processes and that its failure modes involve over-reporting of differing rate constants k_i_ ≠ k_j_ rather than over-reporting of identical rate constants k_i_ = k_j_.

### Hypoexponential fitting suggests an “asymmetric” model for ACE2 action in SARS-CoV-2 fusion

We next applied hypoexponential fitting to help understand the rate-limiting steps in SARS-CoV-2 fusion kinetics. Briefly, SARS-CoV-2 entry involves spike protein binding to cellular ACE2 receptors(34,35) and proteolytic activation of the spike to trigger fusion (36,37). ACE2 facilitates a conformational change in the spike protein (38) that is linked to fusion. Surprisingly, however, single-virus fusion experiments found that ACE2 is not strictly required for SARS-CoV-2 fusion (23). This work measured fusion kinetics with and without ACE2, analyzing the data with fitted gamma distributions. This approach suggested that there were two rate-limiting steps in the absence of ACE2 and one in the presence of ACE2 but did not further illuminate the chemical nature of this difference. Here, we fit hypoexponential functions to the previously measured single-virus fusion kinetics to generate testable mechanistic hypotheses regarding the rate-limiting steps in SARS-CoV-2 entry.

Hypoexponential analysis agrees with the fundamental finding from gamma analysis and provides additional information on SARS-CoV-2’s mechanism of entry. Both approaches predict the most-likely model to be one kinetic step with ACE2 and two kinetic steps without. Interestingly, the rates predicted from hypoexponential fitting where k_1_ and k_2_ can vary freely were 0.0031 and 0.0033 s^-1^ without ACE2 and 0.00338 s^-1^ with ACE2 (Fig. 7). These are highly similar both to each other and to the rates predicted from gamma fitting where k_1_ and k_2_ are constrained to be equal: 0.0030 s^-1^ either with or without ACE2.

**Figure 7.**
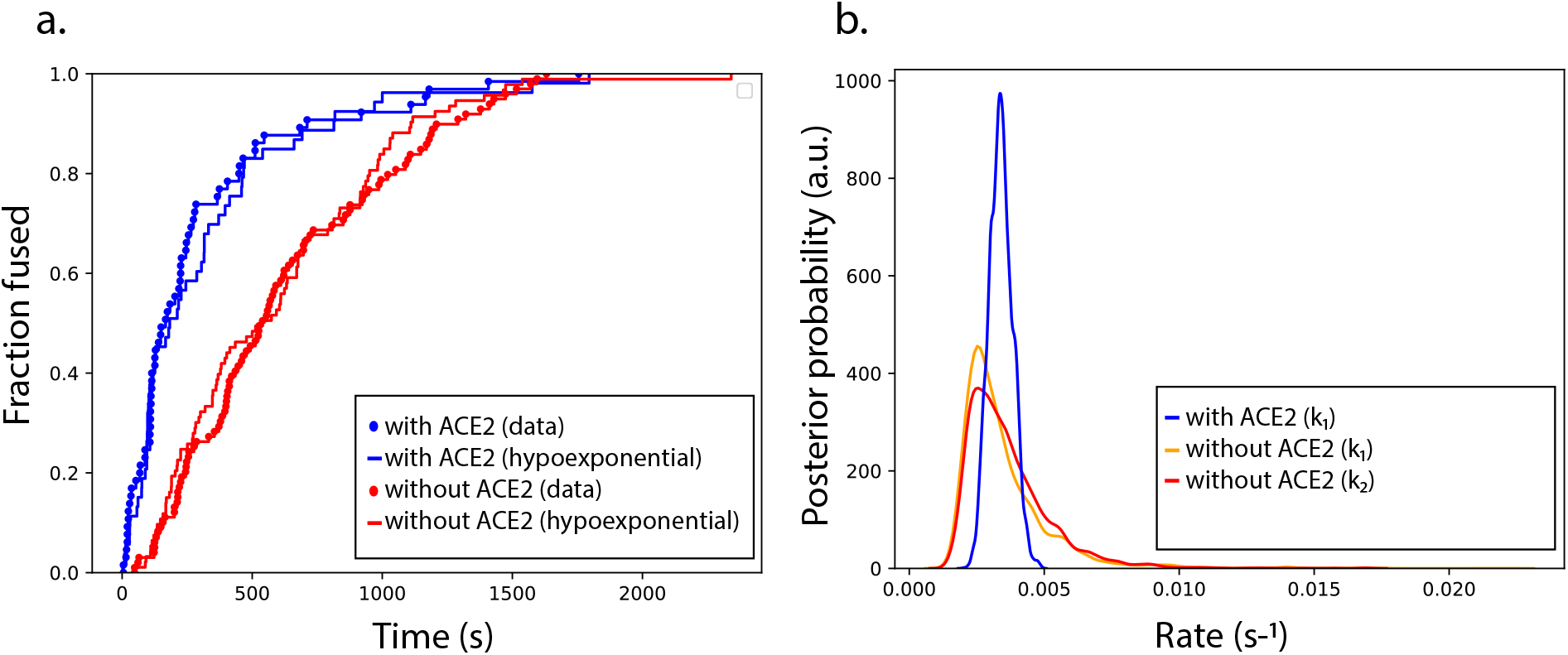
Kinetics of SARS-CoV-2 entry with and without ACE2. Empirical CDFs and hypoexponential fits are plotted in (a), while posterior probability density estimates for the rates are plotted in (b). Maximum-likelihood hypoexponential parameter fits are with ACE2: k=0.00338, N=1 and without ACE2: k_1_ = 0.00310, k_2_ = 0.00327, N=2.

Based on this, we propose the following kinetic model for SARS-CoV-2 fusion kinetics (Fig. 8). Similar to models for influenza discussed in our prior work (33), we formulate a variable-stoichiometry model for fusion protein activation, where docked, unfused virus has an activation free energy for fusion of ΔG_F_^‡^. Each viral protein activation event occurs with a rate k_S_ and lowers the activation barrier to fusion by ΔG_S_, with the corresponding k_F*i*_ rate constants as schematized in the figure. The relative probability of outbound transitions (and thus relative flux) from any given intermediate is thus given by the ratio of first-order rate constants. Hypoexponential fitting can reliably recover rates that differ by less than ∼100x in a three-step process. Therefore, estimation of SARS-CoV-2 fusion as a two-step process in the absence of ACE2 means that either 1) the maximum-flux pathway from U to F in this model is U->US_1_->F or 2) that it is U->US_1_->US_2_->F **and** k_F2_ / k_S_ ≥ 100. Since we estimate the two steps to have near-identical rates in the absence of ACE2, U->US_1_->F is unlikely because kS ∼ kF would an approximately 1:1 mixture of flux through the U->US_1_->F pathway and the U->US_1_->US_2_->F pathway. We therefore conclude the most likely pathway is U->US_1_->US_2_->F with k_F2_ / k_S_ ≥ 100. This implies that ΔG_F_^‡^ = 2ΔG_S_ + *δ*, where *δ* < ΔG_S_, so that k_F2_ >> k_S_ and k_F1_ < k_S_.

**Figure 8.**
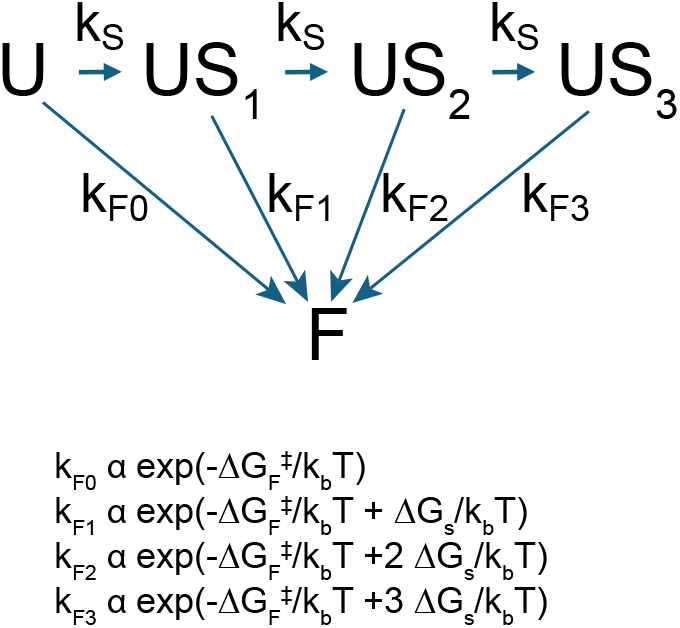
Kinetic scheme for fusion. U, US_n_, and F represent states in the scheme. ΔG_F_^‡^ is the activation energy for fusion, ΔG_s_ is the free energy contributed by each activated spike towards fusion, and k_b_ is Boltzmann’s constant, and T is temperature.

We then consider the kinetic data in the presence of ACE2. Structural and smFRET data suggest that ACE2 affects the activation of SARS-CoV-2 spikes but would be unlikely to alter ΔG_F_ (36,38-40), the energetic contribution of these spikes to fusion. Therefore, our favored hypothesis is that ACE2 affects k_S_ nonuniformly: the most probable pathway in our scenario is that one spike activation event is ACE2-accelerated and not detected as rate-limiting, whereas one spike activation event is not ACE2-accelerated and remains at the original rate k_S_.

This seemingly counterintuitive “asymmetric” scenario could result from two key properties of ACE2-spike interactions: it may be that the number of ACE2 in the viral contact region is typically low and thus the encounter between a second spike and ACE2 may be slow compared to spontaneous activation of that spike. Alternatively, ACE2 has been observed to drive spontaneous inactivation of spike trimers that do not immediately proceed to fusion (23). This inactivation process would limit the time window for recruitment of a second spike, making it unlikely that two spikes remain active during the same time window. We propose future experiments to resolve this by measuring SARS-CoV-2 fusion kinetics as a function of ACE2 membrane density. Such experiments should further constrain mechanistic models in combination with experimentally derived estimates of ACE2 inactivation timescales.

## Discussion

In any biomolecular process, the rate-limiting steps could be singular, multiple and identical, or multiple and heterogeneous. When available data consist of single-molecule dwell times (reporting only on reaction completion), pre-existing approaches often differentiate poorly between the last two cases. We have shown that hypoexponential fits can reliably extract heterogeneous rate constants over a 2-3 order of magnitude range. Since this corresponds to approximately 7-10 k_b_T, this permits a reasonable capability to probe different mechanistic steps. Even if inclusion of heterogeneous rates at the high end of this detection range contributes only slightly to recovering the measured dwell-time distribution, their detection can have important mechanistic consequences.

In this work, we demonstrate a case where definitively ruling out heterogeneous rates within this range yields biological insight. Prior analysis of the role of ACE2 in promoting SARS-CoV-2 fusion had identified a change in the number of rate-limiting steps but could not differentiate whether the component steps were all identical or involved different mechanistic processes. The ability to detect heterogeneous rates, and in this case, the uniformity of rates detected, helps us assign all the identical rate-limiting steps to ACE2-mediated SARS-CoV-2 spike activation. This yields a novel and experimentally testable hypothesis: that SARS-CoV-2 fusion involves one ACE2-accelerated activation event and one ACE2-independent activation event. This seemingly counterintuitive prediction could result if the likelihood of two spikes being simultaneously activated by ACE2 molecules were small and if the rate of encounter between a second spike and a second ACE2 is slower than the rate of ACE2-independent activation (estimated here at 0.003 s^-1^). Such an event could also occur if one spike undergoes spontaneous activation prior to ACE2 encounter by the second spike. This can be tested and compared to physiological virus-cell encounters by experimentally tuning the virus-ACE2 encounter rate.

## Acknowledgements

The authors thank Robert Rawle for helpful discussions and influenza single-event dwell-time data. Marcos Cervantes provided single-event dwell-time data for SARS-CoV-2 fusion. This work was supported by NIGMS R01GM138444 to P.M.K. Work in Sweden was supported by the Knut and Alice Wallenberg Foundation KAW 2020.0209. O.A. was also supported by a traineeship under NSF 2021791.

## Notes

### Competing Interest Statement

The authors have declared no competing interest.

## References

1. Kou, S. C., B. J. Cherayil, W. Min, B. P. English, and X. S. Xie. 2005. Single-molecule Michaelis-Menten equations. J Phys Chem B. 109(41):19068–19081, doi: 10.1021/jp051490q, https://www.ncbi.nlm.nih.gov/pubmed/16853459.

2. van de Meent, J. W., J. E. Bronson, C. H. Wiggins, and R. L. Gonzalez, Jr. 2014. Empirical Bayes methods enable advanced population-level analyses of single-molecule FRET experiments. Biophys J. 106(6):1327–1337, doi: 10.1016/j.bpj.2013.12.055, https://www.ncbi.nlm.nih.gov/pubmed/24655508.

3. Mullner, F. E., S. Syed, P. R. Selvin, and F. J. Sigworth. 2010. Improved hidden Markov models for molecular motors, part 1: basic theory. Biophys J. 99(11):3684–3695, doi: 10.1016/j.bpj.2010.09.067, http://www.ncbi.nlm.nih.gov/pubmed/21112293.

4. Sgouralis, I., and S. Presse. 2017. An Introduction to Infinite HMMs for Single-Molecule Data Analysis. Biophys J. 112(10):2021–2029, doi: 10.1016/j.bpj.2017.04.027, https://www.ncbi.nlm.nih.gov/pubmed/28538142.

5. Fox, E. B., E. B. Sudderth, M. I. Jordan, and A. S. Willsky (2008). An HDP-HMM for systems with state persistence.

6. Syed, S., F. E. Mullner, P. R. Selvin, and F. J. Sigworth. 2010. Improved hidden Markov models for molecular motors, part 2: extensions and application to experimental data. Biophys J. 99(11):3696–3703, doi: 10.1016/j.bpj.2010.09.066, http://www.ncbi.nlm.nih.gov/pubmed/21112294.

7. Teh, Y. W., M. I. Jordan, M. J. Beal, and D. M. Blei. 2006. Hierarchical Dirichlet Processes. Journal of the American Statistical Association. 101(476):1566–1581, http://www.jstor.org/stable/27639773.

8. Siekmann, I., L. E. Wagner, 2nd, D. Yule, C. Fox, D. Bryant, E. J. Crampin, and J. Sneyd. 2011. MCMC estimation of Markov models for ion channels. Biophys J. 100(8):1919–1929, doi: 10.1016/j.bpj.2011.02.059, https://www.ncbi.nlm.nih.gov/pubmed/21504728.

9. Munro, J. B., J. Gorman, X. Ma, Z. Zhou, J. Arthos, D. R. Burton, W. C. Koff, J. R. Courter, A. B. Smith, 3rd, P. D. Kwong, S. C. Blanchard, and W. Mothes. 2014. Conformational dynamics of single HIV-1 envelope trimers on the surface of native virions. Science. 346(6210):759–763, doi: 10.1126/science.1254426, https://www.ncbi.nlm.nih.gov/pubmed/25298114.

10. Sgouralis, I., S. Madaan, F. Djutanta, R. Kha, R. F. Hariadi, and S. Presse. 2019. A Bayesian Nonparametric Approach to Single Molecule Forster Resonance Energy Transfer. J Phys Chem B. 123(3):675–688, doi: 10.1021/acs.jpcb.8b09752, https://www.ncbi.nlm.nih.gov/pubmed/30571128.

11. McKinney, S. A., C. Joo, and T. Ha. 2006. Analysis of single-molecule FRET trajectories using hidden Markov modeling. Biophys J. 91(5):1941–1951, doi: 10.1529/biophysj.106.082487.

12. Floyd, D., J. R. Ragains, J. J. Skehel, S. C. Harrison, and A. M. van Oijen. 2008. Single-particle kinetics of influenza virus membrane fusion. Proc Natl Acad Sci U S A. 105(40):15382–15387, 2556630, .

13. Sarkar, S. K., B. Marmer, G. Goldberg, and K. C. Neuman. 2012. Single-molecule tracking of collagenase on native type I collagen fibrils reveals degradation mechanism. Curr Biol. 22(12):1047–1056, doi: 10.1016/j.cub.2012.04.012, https://www.ncbi.nlm.nih.gov/pubmed/22578418.

14. Ivanovic, T., J. L. Choi, S. P. Whelan, A. M. van Oijen, and S. C. Harrison. 2013. Influenza-virus membrane fusion by cooperative fold-back of stochastically induced hemagglutinin intermediates. Elife. 2:e00333, doi: 10.7554/eLife.00333, http://www.ncbi.nlm.nih.gov/entrez/query.fcgi?cmd=Retrieve&db=PubMed&dopt=Citation&list_uids=23550179.

15. Ivanovic, T., and S. C. Harrison. 2015. Distinct functional determinants of influenza hemagglutinin-mediated membrane fusion. Elife. 4:e11009, doi: 10.7554/eLife.11009, http://www.ncbi.nlm.nih.gov/pubmed/26613408.

16. Chao, L. H., D. E. Klein, A. G. Schmidt, J. M. Pena, and S. C. Harrison. 2014. Sequential conformational rearrangements in flavivirus membrane fusion. Elife. 3:e04389, doi: 10.7554/eLife.04389, http://www.ncbi.nlm.nih.gov/pubmed/25479384.

17. Rawle, R. J., E. R. Webster, M. Jelen, P. M. Kasson, and S. G. Boxer. 2018. pH dependence of Zika membrane fusion kinetics reveals an off-pathway state. ACS Central Science. 4(11):1503–1510.

18. Weber, T. S., I. Jaehnert, C. Schichor, M. Or-Guil, and J. Carneiro. 2014. Quantifying the length and variance of the eukaryotic cell cycle phases by a stochastic model and dual nucleoside pulse labelling. PLoS Comput Biol. 10(7):e1003616, doi: 10.1371/journal.pcbi.1003616, https://www.ncbi.nlm.nih.gov/pubmed/25058870.

19. Yates, C. A., M. J. Ford, and R. L. Mort. 2017. A Multi-stage Representation of Cell Proliferation as a Markov Process. Bulletin of Mathematical Biology. 79(12):2905–2928, doi: 10.1007/s11538-017-0356-4, https://doi.org/10.1007/s11538-017-0356-4.

20. Akaike, H. 1974. A new look at the statistical model identification. Automatic Control, IEEE Transactions on. 19(6):716–723.

21. Sugiura, N. 1978. Further analysis of the data by akaike’s information criterion and the finite corrections: Further analysis of the data by akaike’s. Communications in Statistics-theory and Methods. 7(1):13–26.

22. Rawle, R. J., A. M. Villamil Giraldo, S. G. Boxer, and P. M. Kasson. 2019. Detecting and Controlling Dye Effects in Single-Virus Fusion Experiments. Biophys J. 117(3):445–452, doi: 10.1016/j.bpj.2019.06.022, https://www.ncbi.nlm.nih.gov/pubmed/31326109.

23. Cervantes, M., T. Hess, G. G. Morbioli, A. Sengar, and P. M. Kasson. 2023. The ACE-2 receptor accelerates but is not biochemically required for SARS-CoV-2 membrane fusion. Chemical Science. 14:6997–7004, PMC10306070, doi: 10.1039/D2SC06967A, http://biorxiv.org/content/early/2022/10/24/2022.10.22.513347.abstract.

24. Gillespie, D. T. 1976. A general method for numerically simulating the stochastic time evolution of coupled chemical reactions. Journal of Computational Physics. 22(4):403–434, doi: 10.1016/0021-9991(76)90041-3, https://www.sciencedirect.com/science/article/pii/0021999176900413.

25. Gillespie, D. T. 2007. Stochastic simulation of chemical kinetics. Annu Rev Phys Chem. 58:35–55, doi: 10.1146/annurev.physchem.58.032806.104637, https://www.ncbi.nlm.nih.gov/pubmed/17037977.

26. Rawle, R. J., S. G. Boxer, and P. M. Kasson. 2016. Disentangling Viral Membrane Fusion from Receptor Binding Using Synthetic DNA-Lipid Conjugates. Biophys J. 111(1):123–131, doi: 10.1016/j.bpj.2016.05.048, https://www.ncbi.nlm.nih.gov/pubmed/27410740.

27. Cervantes, M., S. Mannsverk, T. Hess, D. Filipe, A. Villamil Giraldo, and P. M. Kasson. 2025. Single-Virus Microscopy of Biochemical Events in Viral Entry. JACS Au. 5(1):399–407, doi: 10.1021/jacsau.4c00992, https://www.ncbi.nlm.nih.gov/pubmed/39886585.

28. Shaevitz, J. W., S. M. Block, and M. J. Schnitzer. 2005. Statistical kinetics of macromolecular dynamics. Biophys J. 89(4):2277–2285, doi: 10.1529/biophysj.105.064295, https://www.ncbi.nlm.nih.gov/pubmed/16040752.

29. Pabis, A., R. J. Rawle, and P. M. Kasson. 2020. Influenza hemagglutinin drives viral entry via two sequential intramembrane mechanisms. Proc Natl Acad Sci U S A. 117(13):7200–7207, doi: 10.1073/pnas.1914188117, https://www.ncbi.nlm.nih.gov/pubmed/32188780.

30. Wessels, L., M. W. Elting, D. Scimeca, and K. Weninger. 2007. Rapid membrane fusion of individual virus particles with supported lipid bilayers. Biophys J. 93(2):526–538, http://www.ncbi.nlm.nih.gov/entrez/query.fcgi?cmd=Retrieve&db=PubMed&dopt=Citation&list_uids=17449662

31. Li, T., Z. Li, E. E. Deans, E. Mittler, M. Liu, K. Chandran, and T. Ivanovic. 2021. The shape of pleomorphic virions determines resistance to cell-entry pressure. Nature Microbiology. 6(5):617–629, doi: 10.1038/s41564-021-00877-0, https://doi.org/10.1038/s41564-021-00877-0.

32. Floyd, D. L., S. C. Harrison, and A. M. van Oijen. 2010. Analysis of kinetic intermediates in single-particle dwell-time distributions. Biophys J. 99(2):360–366, doi: 10.1016/j.bpj.2010.04.049, https://www.ncbi.nlm.nih.gov/pubmed/20643053.

33. Villamil Giraldo, A. M., and P. Kasson. 2020. Bilayer-Coated Nanoparticles Reveal How Influenza Viral Entry Depends on Membrane Deformability but Not Curvature. Journal of Physical Chemistry Letters. 11:7190–7196, doi: 10.1021/acs.jpclett.0c01778.

34. Zhou, P., X. L. Yang, X. G. Wang, B. Hu, L. Zhang, W. Zhang, H. R. Si, Y. Zhu, B. Li, C. L. Huang, H. D. Chen, J. Chen, Y. Luo, H. Guo, R. D. Jiang, M. Q. Liu, Y. Chen, X. R. Shen, X. Wang, X. S. Zheng, K. Zhao, Q. J. Chen, F. Deng, L. L. Liu, B. Yan, F. X. Zhan, Y. Y. Wang, G. F. Xiao, and Z. L. Shi. 2020. A pneumonia outbreak associated with a new coronavirus of probable bat origin. Nature. doi: 10.1038/s41586-020-2012-7, https://www.ncbi.nlm.nih.gov/pubmed/32015507.

35. Hoffmann, M., H. Kleine-Weber, S. Schroeder, N. Kruger, T. Herrler, S. Erichsen, T. S. Schiergens, G. Herrler, N. H. Wu, A. Nitsche, M. A. Muller, C. Drosten, and S. Pohlmann. 2020. SARS-CoV-2 Cell Entry Depends on ACE2 and TMPRSS2 and Is Blocked by a Clinically Proven Protease Inhibitor. Cell. 181(2):271–280 e278, doi: 10.1016/j.cell.2020.02.052, https://www.ncbi.nlm.nih.gov/pubmed/32142651.

36. Benton, D. J., A. G. Wrobel, P. Xu, C. Roustan, S. R. Martin, P. B. Rosenthal, J. J. Skehel, and S. J. Gamblin. 2020. Receptor binding and priming of the spike protein of SARS-CoV-2 for membrane fusion. Nature. 588(7837):327–330, doi: 10.1038/s41586-020-2772-0, https://doi.org/10.1038/s41586-020-2772-0.

37. Koch, J., Z. M. Uckeley, P. Doldan, M. Stanifer, S. Boulant, and P.-Y. Lozach. 2021. TMPRSS2 expression dictates the entry route used by SARS-CoV-2 to infect host cells. The EMBO journal. 40(16):e107821, doi: 10.15252/embj.2021107821, https://doi.org/10.15252/embj.2021107821.

38. Lu, M., P. D. Uchil, W. Li, D. Zheng, D. S. Terry, J. Gorman, W. Shi, B. Zhang, T. Zhou, S. Ding, R. Gasser, J. Prévost, G. Beaudoin-Bussières, S. P. Anand, A. Laumaea, J. Grover, L. Liu, D. D. Ho, J. R. Mascola, A. Finzi, P. D. Kwong, J. S. Blanchard, and W. Mothes. 2020. Real-time conformational dynamics of SARS-CoV-2 spikes on virus particles. Cell host & microbe. 28(6):880–891.

39. Shang, J., G. Ye, K. Shi, Y. Wan, C. Luo, H. Aihara, Q. Geng, A. Auerbach, and F. Li. 2020. Structural basis of receptor recognition by SARS-CoV-2. Nature. doi: 10.1038/s41586-020-2179-y, https://www.ncbi.nlm.nih.gov/pubmed/32225175.

40. Yan, R., Y. Zhang, Y. Li, L. Xia, Y. Guo, and Q. Zhou. 2020. Structural basis for the recognition of SARS-CoV-2 by full-length human ACE2. Science. 367(6485):1444–1448, doi: 10.1126/science.abb2762, https://www.ncbi.nlm.nih.gov/pubmed/32132184.

